# RNA-sequencing analysis of a multistep and hit-and-run cell and animal model of KSHV tumorigenesis reveal the roles of mutations, CpG methylation, and viral-infection footprints in oncogenesis

**DOI:** 10.1101/792028

**Authors:** Julian Naipauer, Daria Salyakina, Guy Journo, Santas Rosario, Sion Williams, Martin Abba, Meir Shamay, Enrique A. Mesri

## Abstract

Human viral oncogenesis is the consequence of cell transformation mediated by virally encoded oncogenes in combination with host oncogenic alterations. Kaposi’s sarcoma (KS), caused by the Kaposi’s sarcoma-associated herpes virus (KSHV), is an AIDS-associated cancer characterized by angiogenesis and spindle-cells proliferation. KSHV-infected KS lesions are composed of latently-infected cells, as well as cells expressing lytic genes that have been implicated in the development of the KS angioproliferative phenotype. The existence of KS lesions with varying levels of KSHV-infected cells suggests also the existence of virus-independent “hit-and-run” mechanisms of sarcomagenesis, whereby viral infection irreversibly induce genetic or epigenetic oncogenic alterations in host cells. We have integrated genetic mutations, changes in expression signatures and methylation analysis to dissect genetic and epigenetic signaling pathways in an unbiased manner in the mECK36 mouse model of KSHV tumorigenesis. Pathway analysis of differential expressed genes (DEGs) showed KSHV lytic switch, DNA methylation and Epigenetic as the most regulated pathways during KSHV-dependent *in vivo* tumorigenesis. Methylation analysis data indicates that during the development of KSHV-infected tumors the most changes were towards hypo-methylation of tissues specific genes and oncogenic signature pathways, on the other hand during viral loss and development of KSHV-negative tumors changes are towards hyper-methylation. Mutational analysis of KSHV-infected cells and tumors revealed a set of mutations, including mutations in three inflammasome-related IFN response genes, that were absent in KSHV-infected cells but present in all KSHV-infected tumors in the same loci pointing to clonal selection “in vivo”. This result suggests that in the context of *in vivo* tumorigenesis both these mutations and the virus may determine tumor growth. On the other hand, clustering analysis of mutations driving KSHV-negative tumors reveal a network comprising PDGFRA D842V, Pak1 and Nucleolin mutations implicated in cell proliferation. Our results have uncovered novel specific aspects of the interplay between host oncogenic alterations and virus-induced transcriptional effects as well as the epigenetic changes induced by KSHV infection and tumorigenesis. The existence virally-induced irreversible genetic and epigenetic oncogenic alterations support the possibility for hit-and-run KSHV sarcomagenesis which is consistent with pathological and clinical findings.

**AUTHOR SUMMARY:** We performed whole genome RNA sequencing and CpG DNA methylation analysis in a mouse bone-marrow endothelial-lineage cells (mEC) transfected with the KSHVBac36 (mECK36 cells), that are able to form KSHV-infected tumors in nude mice, which were thoroughly characterized as KS-like tumors. This unique model allowed us to dissect genetic and epigenetic mechanisms of KSHV dependent and hit-and-run sarcomagenesis. We found that during KSHV *in vivo* lytic switch and KSHV-dependent tumorigenesis DNA methylation and Epigenetic regulation are among the most host-regulated pathways. CpG DNA methylation analysis during transformation supports the notion that loss of methylation (hypo-methylation) is the major epigenetic change during this process. Sequence analysis of KSHV-positive tumors revealed that KSHV tumorigenesis not only selects for the presence of the virus but also pre-existing host mutations that allow the KSHV oncovirus to express the oncogenic lytic program and creates a permissive environment of inflammation and viral tumorigenesis providing a selective advantage *in vivo*.

## INTRODUCTION

Human viral oncogenesis is the consequence of the transforming activity of virally encoded oncogenes in combination with host oncogenic alterations (Mesri et al., 2014). Kaposi’s sarcoma (KS), caused by the Kaposi’s sarcoma-associated herpes virus (KSHV), is a major cancer associated with AIDS and is consequently a major global health challenge (Cesarman et al., 2019; Dittmer and Damania, 2016; Mesri et al., 2010). The KS tumors are characterized by intense angiogenesis and the proliferation of spindle cells that can affect the skin, mucosa and viscera, causing significant morbidity (Cesarman et al., 2019; Dittmer and Damania, 2016; Mesri et al., 2010). Although KS can be treated with anti-retroviral therapy and chemotherapy, it is estimated that more than a half of AIDS associated KS patients will not be cured (Cavallin et al., 2014; Cesarman et al., 2019). Understanding the interplay between viral and cellular genes leading to KS carcinogenesis is paramount to developing rationally designed therapies for KS.

KSHV-infected KS lesions are composed of a majority of latently infected cells, as well as cells expressing lytic genes that have been implicated in the development of the KS angioproliferative phenotype via paracrine and autocrine mechanisms (Bais et al., 1998; Cesarman et al., 2000; Dittmer and Damania, 2016; Ganem, 2010; Mesri et al., 2010; Montaner et al., 2006). Canonical latent and lytic KSHV infections are not highly oncogenic as suggested by the low incidence of KS in the general KSHV-seropositive population, the low efficiency cell transformation in vitro(Flore et al., 1998), and the inability of KSHV latent infection per se to transform endothelial cells (Lagunoff et al., 2002). Thus, a critical question becomes how the interplay between KSHV and host gene expression leads to cell transformation and establishment of the KS angio-proliferative lesion. We proposed a mechanism for *vGPCR* immortalization of ECs involving KDR autocrine activation compatible with such a modality of hit and run oncogenicity (Bais et al., 2003). A similar mechanism of hit and run autocrine immortalization has been proposed for *HTLV-1 Tax*, which immortalizes T cells by creation of an IL-2/IL-2R autocrine loop, but is deleted from the provirus in HTLV-1-transformed IL-2-dependent adult T cell lymphoma cells (Cesarman and Mesri, 1999). We have developed a cell and animal model based on mouse bone marrow cells of the endothelial cell lineage transfected with a recombinant KSHV genome (mECK36 tumor model) (Mutlu et al., 2007). mECK36 tumors show consistent expression of the KS markers such as Podoplanin and LYVE-1. This expression occurs with concomitant up-regulation of KSHV lytic oncogenes and angiogenesis ligands/receptors, pointing to the upregulation of various receptor tyrosine kinase signaling axes (Mutlu et al., 2007). Using this model we showed that the most prominently activated tyrosine kinase was PDGFRA (Cavallin et al., 2018), which was activated by lytic KSHV genes such as vGPCR via a ligand-mediated mechanism and necessary for KSHV sarcomagenesis (Cavallin et al., 2018). More importantly, PDGFRA was prominently expressed and phosphorylated in the vast majority of AIDS-KS tumors (Cavallin et al., 2018). Yet, we were able to find a few biopsies in which only a small percentage of phospho-PDGFRA+ve tumor spindle cells were infected with KSHV as evidenced by KSHV latent antigen LANA staining. The existence of KS lesions with varying levels of KSHV infected (LANA+) cells suggests the existence of virus-independent hit-and-run mechanisms of sarcomagenesis. In hit-and-run scenario, cells are irreversibly transformed by KSHV and are able to sustain tumors in the absence of virus. This is consistent with some reports that have identified the presence of host oncogenic mutations in KS lesions (Nicolaides et al., 1994; Pillay et al., 1999). Yet the possibility of KSHV induced hit-and-run tumorigenesis and its mechanistic underpinnings have not been yet firmly established.

DNA methylation at CpG dinucleotides is an epigenetic mark that has been studied extensively in the context of cancer. Methylation of the cytosine residue in the CpG dinucleotide is carried out by the DNA methyltransferases DNMT1, DNMT3a, and DNMT3b (Bestor, 2000). Many promoters contain CpG islands, and these islands are protected from methylation in normal tissues (Antequera et al., 1990). In cancer cells, some of these CpG islands become aberrantly hypermethylated, and this is usually correlated with transcription repression (Baylin, 2005). On the other hand, global hypomethylation has been described in cancer cells (Goelz et al., 1985). DNA methylation is regulated by KSHV on several levels. The latency-associated nuclear antigen (LANA/ORF73) encoded by KSHV leads to CpG methylation by interacting with the cellular *de novo* DNA methyltransferase, DNMT3a, and recruiting DNMT3a to certain cellular promoters that become methylated and repressed (Shamay et al., 2006). An additional mechanism by which KSHV might modify the human methylome is via the Polycomb complex that creates the histone mark histone H3 trimethylated on Lys27 (H3K27me3) and can direct cellular CpG methylation via its interaction with DNMTs (Gunther and Grundhoff, 2010; Schlesinger et al., 2007). The pattern of CpG DNA methylation in chronically infected primary effusion lymphoma (PEL) cells and during de-novo in-vitro infection was investigated for the KSHV episomal genome (Chen et al., 2001; Gunther and Grundhoff, 2010; Shamay et al., 2012a) and for the host cellular genome (Journo et al., 2018).

A unique feature of the mECK36 cell model is that KSHV tumorigenesis is tightly linked to the presence of the virus since mECK36 cells that lose the viral episome can survive in culture but are not tumorigenic. Yet, upon mECK36 tumor formation in mice, explanted cells that are forced to lose the viral episome continue being tumorigenic; in part and as we have recently shown, by the irreversible *in vivo* acquisition of host mutations such as PDGFRA D842V (Cavallin et al., 2018). Therefore, the combination of the mECK36 cells and their derivatives *in vitro* and *in vivo* constitute a unique model to study through the use of Next Generation Sequencing not only the transcriptional events regulated by KSHV *in vitro* and *in vivo*; but more interestingly, to define the viral oncogenic footprint at the DNA level encompassing both changes in CpG methylation as well as mutations in transcribed genes.

In this work, we have integrated genetic mutations, changes in expression signatures and methylation analysis during direct and hit-and-run KSHV oncogenesis *in vitro* and *in vivo* to dissect genetic and epigenetic signaling pathways in an unbiased manner in the mECK36 mouse model of KSHV tumorigenesis (Mutlu et al., 2007). Pathway analysis of differential expressed genes (DEGs) during KSHV-dependent *in vivo* tumorigenesis showed KSHV lytic switch and DNA methylation and Epigenetic as the most regulated pathways during this process. Our methylation analysis data indicates that during the development of tumors the most profound changes are towards hypo-methylation of tissues specific genes and oncogenic signature pathways as well as for KSHV genes, on the other hand during viral loss and development of KSHV (−) tumors the most profound changes are towards hyper-methylation of these and additional oncogenic pathways. Mutational analysis of mECK36 KSHV (+) cells and tumors revealed a surprising set of mutations, including mutations in three inflammasome related IFN response genes, absent in KSHV (+) cells but present in all KSHV (+) tumors in the same location. This indicates that these mutations should be the consequence of *in vivo* clonal selection of few mutated KSHV (+) cells of the population. This result suggests that in the context of *in vivo* tumorigenesis both these mutations and the virus may determine tumor growth. On the other hand, clustering analysis of mutations driving KSHV (−) tumors reveal a network comprising PDGFRA D842V, Pak1 and Nucleolin mutations implicated in cell proliferation. Our results have uncovered novel specific aspects of the interplay between host oncogenic alterations and virus-induced transcriptional effects as well as the relationship between epigenetic changes induced by KSHV infection and tumorigenesis. These virally-induced irreversible oncogenic alterations support the possibility for hit-and-run KSHV sarcomagenesis which is consistent with pathological and clinical findings.

## RESULTS

### Animal Model of Multistep and Hit-and-Run KSHV sarcomagenesis

Mouse bone-marrow endothelial-lineage cells (mEC) transfected with the KSHVBac36 (mECK36 cells) are able to form KSHV-infected tumors in nude mice, which were thoroughly characterized as KS-like tumors (Mutlu et al., 2007), thus providing a platform to dissect molecular mechanisms of tumorigenesis by KSHV. When mECK36 cells bearing Bac36KSHV (abbreviated as **KSHV (+) cells** see **Figure 1A**) lose the KSHV episome *in vitro* by withdrawal of antibiotic selection (abbreviated as **KSHV (−) cells**), they completely lose tumorigenicity (Mutlu et al., 2007). Tumors formed by mECK36 cells are all episomally infected with KSHVBac36 (Mutlu et al., 2007)(abbreviated as **KSHV (+) tumors**). Cells explanted from these tumors and grown in absence of antibiotic lose the KSHV episome (abbreviated as **KSHV (−) tumor cells**), are tumorigenic and are able to form KSHV-negative tumors (abbreviated as **KSHV (−) tumors**) indicating that they became irreversibly transformed by KSHV during *in vivo* growth. This is likely due to host genetic and/or epigenetic alterations accumulated during *in vivo* tumor growth that can compensate for KSHV-induced tumorigenicity after loss of the KSHV episomes.. We decided to use this unique animal model of multistep and hit-and-run KSHV sarcomagenesis to dissect transcriptional, genetic and epigenetic mechanisms of KSHV dependent and independent (hit-and-run) sarcomagenesis in an unbiased high-throughput fashion.

**FIGURE 1.**
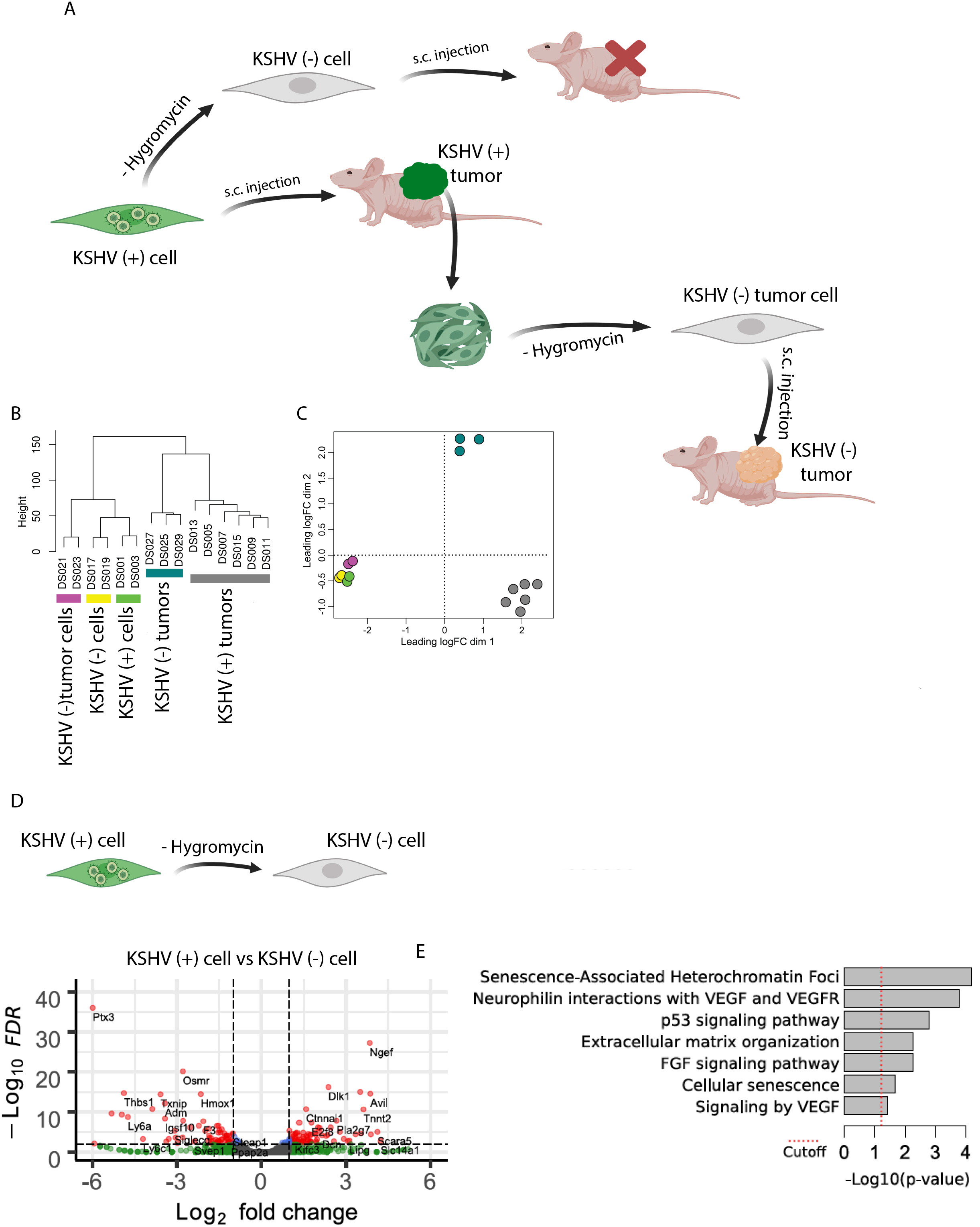
GENOME-WIDE ANALYSIS OF KSHV AND HOST TRANSCRIPTS BY RNA DEEP SEQUENCING. **(A)** Animal Model of Multistep KS Carcinogenesis. **(B)** Hierarchical clustering. The dendrogram of all RNA-Seq samples is shown with the. **(C)** Multidimensional scaling plot KSHV (+) cells versus KSHV (−) cells. **(D)** Scheme of the comparison KSHV (+) cell versus KSHV (−) cell. **(E)** Volcano plot showing 144 differentially expressed genes (DEGs) analyzed by RNA-Seq between KSHV (+) cells and KSHV (−) cells *in vitro*. **(F)** Functional enrichment analysis based on genes differentially expressed among KSHV (+) cells and KSHV (−) cells *in vitro*.

### RNA sequencing analysis of mECK36 model of KSHV tumorigenesis

To identify changes at the transcriptional level in our murine model of KSHV-infected cells and tumors (Ma et al., 2013; Mutlu et al., 2007), high throughput RNA sequencing was performed to identify differences in the gene expression profile, allowing us to perform key biological comparisons (Table 1). We have performed Illumina, stranded, RNA sequencing analysis of all KSHV stages of this cell and animal model including **KSHV (+) cells, KSHV (−) cells, KSHV (−) tumor cells, KSHV (+) tumors and KSHV (−) tumors**. Expression of 16,016 cellular genes in all replicates were identified. Unsupervised clustering (Figure 1B) and Multidimensional scaling plot (Figure 1C) shows how KSHV status and tissue type cluster with each other. Interestingly, cells *in vitro* cluster together whether they are KSHV (+), KSHV (−) or KSHV (−) tumor cells. On the other hand, KSHV (+) and KSHV (−) tumors *in vivo* are separate from each other and from the cells *in vitro* (Figure 1C), further indicating that processes occurring *in vivo* are predominantly distinct from *in vitro* and are distinctly affected by the uniqueness of KSHV-induced tumorigenesis.

**TABLE 1.**
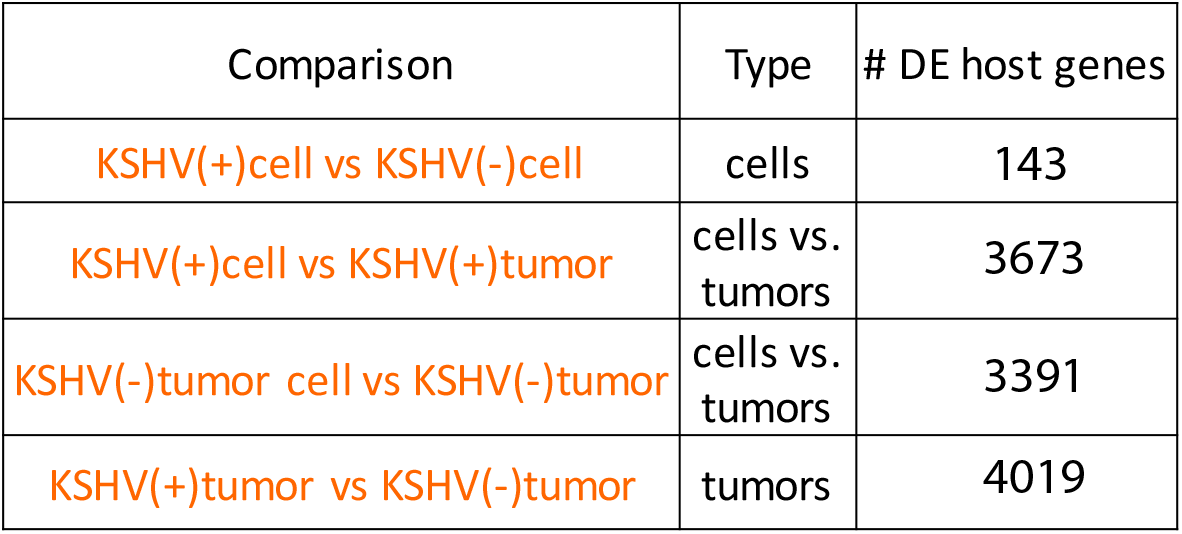
NUMBER OF IDENTIFIED MUTATIONS IN SAMPLES.

To understand the effect of losing the virus *in vitro* and the ability to form tumors *in vivo* of KSHV (−) cells, we compared host gene expression profiles of KSHV (+) cells with KSHV (−) cells (Figure 1D). As expected by the minimal phenotypic *in vitro* differences already described in our previous work (Mutlu et al., 2007), the comparison between KSHV (+) and KSHV (−) cells showed only 143 differentially expressed genes (DEG) (Table 1). Pathway analysis of these DEGs showed changes on Cellular senescence, VEGF signaling (Mutlu et al., 2007), FGF signaling and p53 signaling. The small number of host DEG between tumorigenic KSHV (+) cells and non-tumorigenic KSHV (−) cells highlights the importance of the *in vivo* KSHV lytic switch and environmental cues during the process of *in vivo* tumor formation that was formerly characterized as “ in vivo-restricted oncogenesis” (Mutlu et al., 2007).

To deepen our understanding of the mechanism of KSHV mediated oncogenesis *in vivo* (Figure 2A), we analyzed the KSHV transcriptome using RNA-Sequencing analysis. Multidimensional scaling plot and heat map analysis in Figure 2B-C shows that these two stages (*in vitro* versus *in vivo*) bear two different KSHV gene expression profiles. KSHV (+) tumors showed an increased expression of lytic KSHV genes, including several well-characterized viral oncogenes such as vGPCR (ORF74), vIL6 (ORF-K2) and K15 (Figure 2C). To better understand and visualize these differences we performed a genome-wide tiled analysis of KSHV transcripts in KSHV (+) tumors and KSHV (+) cells (Figure 2D). This result corroborates the marked KSHV lytic *in vivo* switch, previously described for this model of KSHV tumorigenesis using real-time qRT-PCR array (Cavallin et al., 2018; Mutlu et al., 2007) throughout the process of mECK36 tumorigenesis.

**FIGURE 2.**
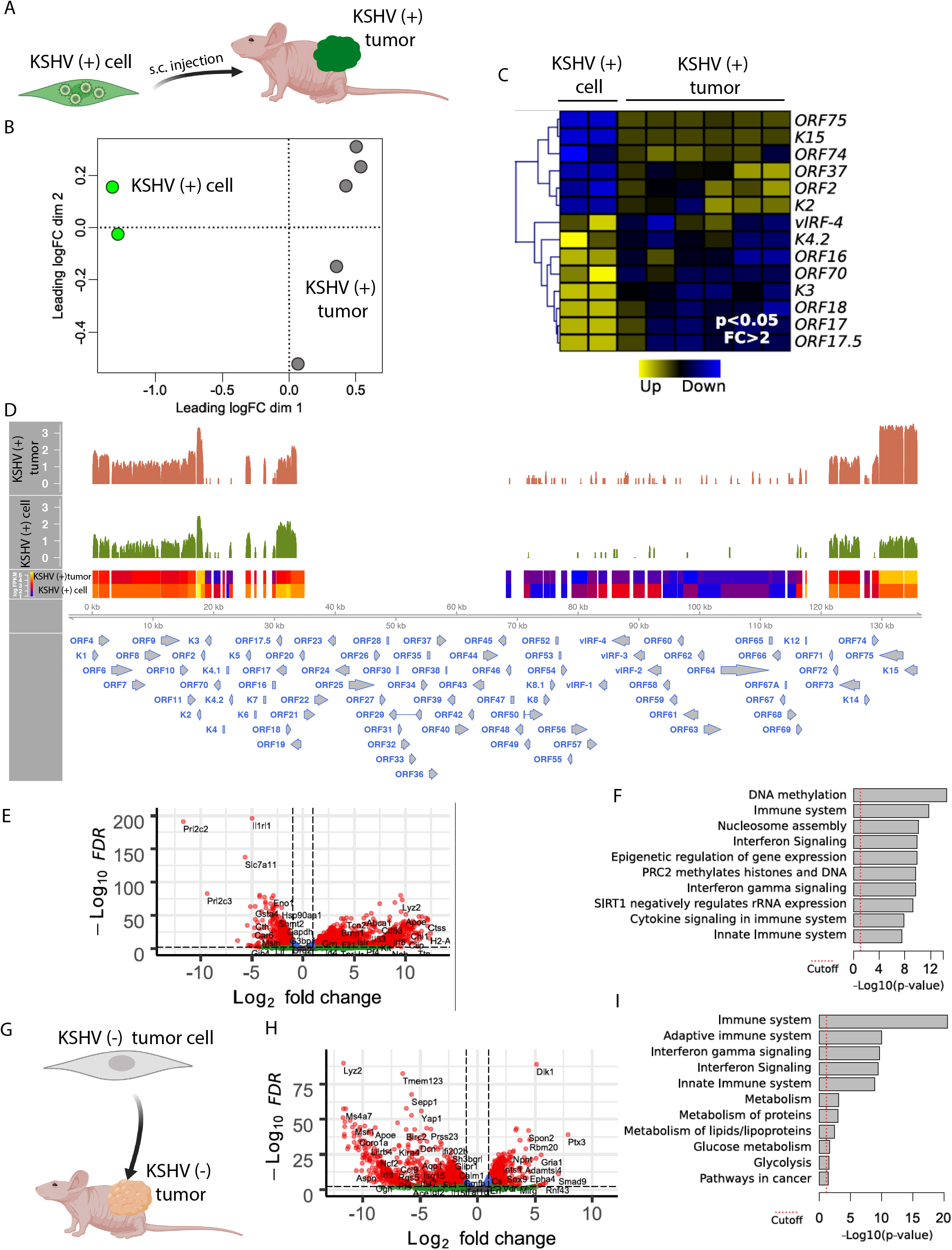
TRANSCRIPTIONAL EFFECTS MEDIATED BY KSHV AND BY ENVIRONMENTAL CUES *IN VIVO*. **(A)** Scheme of the comparison KSHV (+) cell versus KSHV (+) tumor. **(B)** Multidimensional scaling plot for KSHV gene expression of KSHV (+) cells versus KSHV (+) tumors. **(C)** Heat map of KSHV differentially expressed genes between KSHV (+) cells and KSHV (+) tumors. *P < 0.05. **(D)** Genome-wide analysis of KSHV transcripts by RNA deep sequencing, comparison of the transcription profiles of KSHV (+) cells and KSHV (+) tumors. Transcriptional levels of viral genes were quantified in reads per kilobase of coding region per million total read numbers (RPKM) in the sample. The y-axis represents the number of reads aligned to each nucleotide position and x-axis represents the KSHV genome position. **(E)** Volcano plot showing 3674 differentially expressed genes (DEGs) analyzed by RNA-Seq between KSHV (+) cells and KSHV (+) tumors. **(F)** Functional enrichment analysis based on genes differentially expressed among KSHV (+) cells and KSHV (+) tumors. **(G)** Scheme of the comparison KSHV (−) cell versus KSHV (−) tumor. **(H)** Volcano plot showing 3392 differentially expressed genes (DEGs) analyzed by RNA-Seq between KSHV (−) cells and KSHV (−) tumors. **(I)** Functional enrichment analysis based on genes differentially expressed among KSHV (−) cells and KSHV (−) tumors.

To study and compare transcriptional effects in the host genes induced by *in vivo* environmental cues both in KSHV-dependent and KSHV-independent settings, we compared gene expression profiles of KSHV (+) cells versus KSHV (+) tumors (Figure 2E) with that of KSHV (−) tumor cells versus KSHV (−) tumors (Figure 2G). Both comparisons showed around 3000 host DEGs (Table 1 and Figure 2E and 2H), further indicating the impact of *in vivo* growth conditions on host gene expression. Pathway analysis of these DEGs showed DNA methylation and Epigenetic regulation together with Immune System related pathways as the most differentially regulated when KSHV (+) cells form KSHV (+) tumors (Figure 2F). Interestingly, the transition *in vitro* to *in vivo* but in the absence of KSHV, when KSHV (−) tumor cells form KSHV (−) tumors did not show DNA methylation and Epigenetic related regulation pathways as differentially regulated. Instead they showed Immune and Metabolic related pathways as the most differentially regulated pathways (Figure 2I), further indicating the importance of DNA methylation and Epigenetic regulation during KSHV-dependent transformation and tumorigenic growth.

To study the impact of KSHV infection within mECK36 tumors we compared the transcriptome of KSHV (+) tumors with KSHV (−) tumors, which are both driven by PDGFRA signaling (Cavallin et al., 2018). In KSHV infected tumors PDGFRA is activated by KSHV-induced induced ligands (PDGFA and B), while in KSHV (−) tumors PDGFRA bears a heterozygous constitutively activated mutated form (D842V)(Cavallin et al., 2018). Figure 3A shows that gene expression profiles differences in KSHV (+) compared to KSHV (−) tumors ascends to 4019 DEG (Table 1 and Figure 3B), illustrating the impact of *in vivo* KSHV gene expression on host gene expression. Remarkably, pathway analysis showed DNA methylation and Epigenetic regulation of gene expression as the most differentially-regulated pathway between KSHV-dependent and KSHV-independent tumorigenesis (Figure 3C). We used the CIBERSORT *in silico* method (Newman et al., 2015) to determine absolute immune cell fractions within the tumor microenvironment of KSHV-dependent and KSHV-independent tumors. We found the occurrence of Immune cells infiltration (B-cells, neutrophils, NK cells)) in KSHV (+) tumors (Figure 3D) which reflects the contribution of the inflammatory infiltrate to KSHV driven tumors by innate immunity mechanisms that can operate even in athymic mice. Although both KSHV (+) and KSHV (−) tumors are highly vascularized, the CIBERSORT showed also a remarkable increase in endothelial cell component which is consistent to the ability of KSHV to upregulate angiogenesis and endothelial specific genes (Mutlu et al., 2007) and induce transendothelial differentiation (Cheng et al., 2011; Li et al., 2018).

**FIGURE 3.**
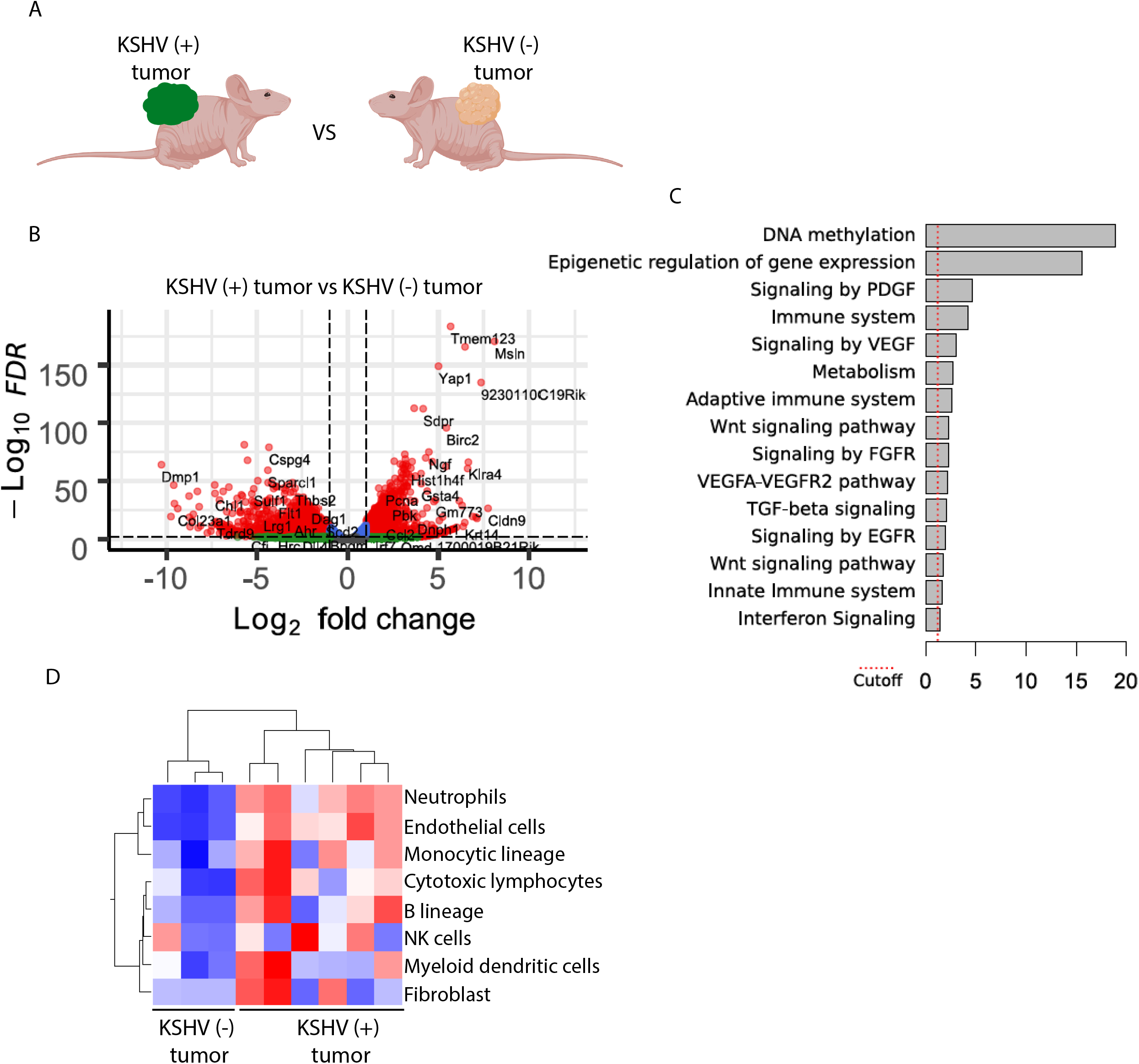
IMPACT OF TRANSCRIPTIONAL EFFECTS OF KSHV INFECTION IN TUMORS. **(A)** Scheme of the comparison KSHV (+) tumors versus KSHV (−) tumors. **(B)** Volcano plot showing 4020 differentially expressed genes (DEGs) analyzed by RNA-Seq between KSHV (+) tumors and KSHV (−) tumors. **(C)** Functional enrichment analysis based on genes differentially expressed among KSHV (+) tumors and KSHV (−) tumors. **(D)** CIBERSORT *in silico* method to determine absolute immune cell fractions within the tumor microenvironment of KSHV-dependent and KSHV-independent tumors.

### Methylation footprint of KSHV infection in the context of KSHV oncogenesis

KSHV is a reprogramming virus that encodes viral genes with powerful epigenetic regulating activities that affect the host. In particular, KSHV could affect DNA CpG methylation on the cellular genome, which contribute to cellular transformation and tumorigenicity. A portion of the DNA CpG methylated sites are expected to remain even after KSHV episomal loss leading to irreversible epigenetic regulatory changes affecting host gene expression. We showed in Figures 2 and 3 the importance of DNA methylation related pathways in the process of KSHV-dependent tumorigenesis. DNA methylation analysis of the cells and tumors generated from this multistep model, affords unique biological comparisons and the possibility of studying the footprints of KSHV infection at the level of the CpG methylation landscape. This afforded remarkable and unique observations on the effects of KSHV infection in the host and some of the molecular mechanisms underpinning KSHV oncogenicity. To follow genome wide DNA methylation, DNA from cells grown in culture or during tumorigenic growth in mice, was subjected to enrichment on Methylated DNA binding beads (MBD2-beads) and the eluted DNA served for library preparation and next-generation-sequencing. The sequenced reads were aligned to the mouse (mm10) genome, and enriched peaks were identified, and annotated. During the transition between KSHV (+) cells and their KSHV (+) tumors (Figure 4A), where we showed a KSHV *in vivo* lytic switch and DNA methylation as the most differentially regulated pathway (Figure 2B-D), we identified 4515 differentially hypo-methylated regions and 3525 differentially hyper-methylated regions (Figure 4B). In order to correlate differential methylation with gene expression we generated a list of differentially methylated promoter (−1500 to +200) regions. Here, clear preferential for hypo-methylation was observed, with 1724 hypo-methylated and 590 hyper-methylated promoters (Figure 4C). Analysis of these hypo-methylated promoter peaks on the GREAT (Genomic Regions Enrichment of Annotations Tool, (http://great.stanford.edu/public/html/index.php) platform identified biological process for pancreatic A cell, astrocyte, dendritic spine, glial cell, enteroendocrine cell, and eye differentiation. In other words, hypo-methylation of many tissue specific promoters, that are expected to be hyper-methylated in the endothelial-lineage cells infected by the virus, and may suggest loss of cell identity/ de-differentiation. These hypo-methylated promoter peaks also identified oncogenic signatures, for genes up-regulated by NF-kB, K-RAS, IL-2, and by knockdown of EED. No terms were identified in biological processes and oncogenic signatures for hyper-methylated promoter peaks (Table S1). Our data indicates that during the development of tumors and KSHV *in vivo* lytic switch the most profound changes are towards hypo-methylation of tissues specific genes and oncogenic signature pathways. Next, we combined our differential promoter methylation data with gene expression data from Figure 2A. This analysis identified 340 hypo-methylated and up-regulated genes, and only 6 hyper-methylated and repressed genes (Figure 4D). Here again, the preference towards hypo-methylation and up-regulation was highlighted. The heatmap of several genes that were both hypo-methylated and up-regulated in KSHV (+) tumors is presented (Figure 4E). These include the two Platelet Derived Growth Factors Pdfga, Pdgfb and their receptor Pdgfra. In a recent study, we have shown that the activation of the PDGF signaling pathway is critical for KS development (Cavallin et al., 2018). We also identified hypo-methylation and up-regulation of Angiopoietin 2 (Angpt2) and 4 (Angpt4) that in combination with VEGF facilitate endothelial cell migration and proliferation, to promote angiogenesis. In addition, hypo-methylation and up-regulation of several neuronal specific, Pax6 and Shank3 genes, and imprinted gene such as Wilms Tumor Protein 1 (WT1), linking KSHV-associated tumorigenesis with loss of cell identity. Chromosome view of three genes that we detected hypo-methylation in the regulatory region, clearly show that the loss of DNA methylation is specific in the gene promoter around the transcription start site (Figure 4F). Gene ontology analysis on STRING (https://string-db.org) for the list of hypo-methylated and up-regulated genes, identified the following cellular pathways; Ras signaling, cytokine receptors, PI3K-Akt signaling, Rap1 signaling, calcium signaling, focal adhesion, and ECM-receptor interaction (Figure 4G). Methylation analysis on KSHV genome showed also a trend to more hyper-methylation in KSHV (+) cells than in KSHV (+) tumors (Figure 4H) correlated with an increase in KSHV viral gene expression in these KSHV (+) tumors (Figure 2 B-D).

**FIGURE 4.**
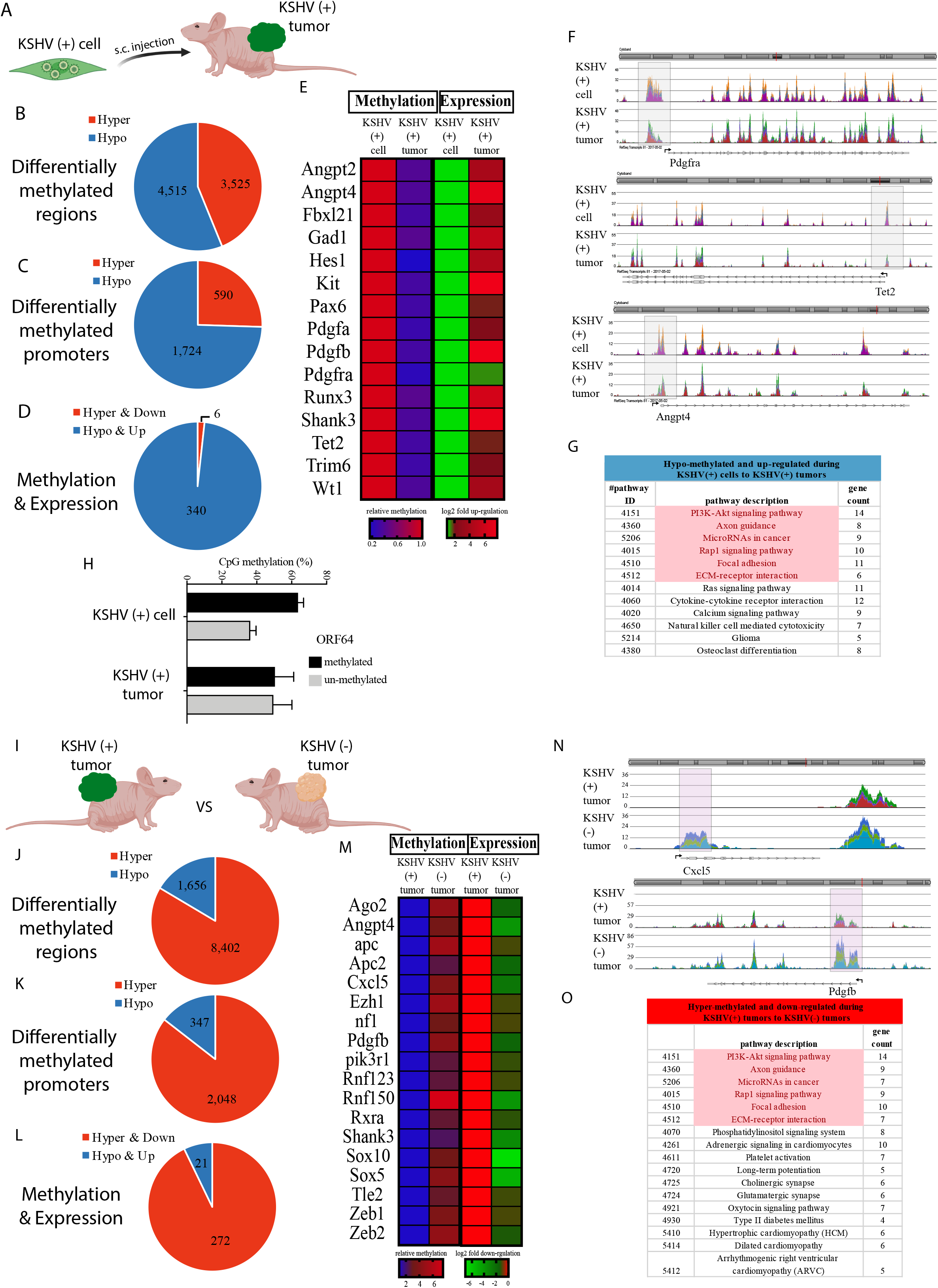
METHYLATION FOOTPRINT OF KSHV INFECTION IN THE CONTEXT OF KSHV ONCOGENESIS. **(A)** Scheme of the comparison KSHV (+) cell versus KSHV (+) tumor. **(B)** The number of differentially methylated regions. Red represents hyper-methylation, and blue represents hypo-methylation. **(C)** Differentially methylated promoters (−1500 to +200 relative to TSS). Red represents hyper-methylation, and blue represents hypo-methylation. **(D)** Differentially methylated promoters that were correlated with gene expression. Red represents hyper-methylation, and blue represents hypo-methylation. **(E)** Heatmap of selected genes with differentially methylated promoters that were correlated with gene expression. **(F)** Chromosome view of two gene promoters which were differentially hypo-methylated. **(G)** Gene ontology analysis on STRING identified pathways that were induced during tumorigenesis with the virus. Common pathways that were repressed following viral loss are marked in pink. **(H)** Evaluation of KSHV methylation was performed on MBD2-beads enriched DNA followed by qPCR of the methylated region in ORF64. Percentage of un-methylated (gray) and methylated (black) fractions are presented. **(I)** Scheme of the comparison KSHV (+) tumors versus KSHV (−) tumors. **(J)** The number of differentially methylated regions. Red represents hyper-methylation, and blue represents hypo-methylation. **(K)** Differentially methylated promoters (−1500 to +200 relative to TSS). Red represents hyper-methylation, and blue represents hypo-methylation. **(L)** Differentially methylated promoters that were correlated with gene expression. Red represents hyper-methylation, and blue represents hypo-methylation. **(M)** Heatmap of selected genes with differentially methylated promoters that were correlated with gene expression. **(N)** Chromosome view of two gene promoters which were differentially hypo-methylated. **(O)** Gene ontology analysis on STRING identified pathways that were repressed during tumorigenesis without the virus. Common pathways that were induced following tumorigenesis with the virus are marked in pink.

Next, we followed the process where KSHV (+) tumors have lost the viral episome by growth without antibiotic selection and re-grown as tumors in mice, KSHV (−) tumors. We identified 8402 differentially hyper-methylated regions and 1656 differentially hypo-methylated regions, with a clear preference for hyper-methylation (Figure 4J). When the list of differential methylation was limited to promoter (−1500 to +200) regions, again a clear preferential for hyper-methylation was observed with 2048 hyper-methylated and only 347 hypo-methylated promoters (Figure 4K). While in the transition between KSHV (+) cells to KSHV (+) tumors hypo-methylation was more significant, in the comparison of KSHV (+) tumors to KSHV (−) tumors hyper-methylation governs. Analysis of these hyper-methylated promoter peaks on the GREAT (Genomic Regions Enrichment of Annotations Tool, (http://great.stanford.edu/public/html/index.php) platform identified biological process for actin filament organization, regulation of cell fate, cell-substrate and cell-matrix adhesion, phagocytosis, epithelial cell differentiation, smooth muscle contraction, and Rac protein signal transduction. The pathways involved in BCR (B-cell receptor) signaling, RXR (retinoid x receptor) and RAR (retinoic acid receptor) signaling. These hyper-methylated promoter peaks also identified oncogenic signatures for SELL, MYD88, RAGE, LTK, and PML. No terms were identified in these categories for hypo-methylated promoter peaks (Table S2). Our data indicates that during KSHV loss and development of KSHV (−) tumors the most profound changes are towards hyper-methylation. When we combined our differential promoter methylation data with gene expression data from Figure 3, we identified, 272 hyper-methylated and down-regulated genes, and only 21 hypo-methylated and up-regulated genes (Figure 4L). Here again, the preference towards hyper-methylation and down-regulation was highlighted. The heatmap of several genes that were both hyper-methylated and down-regulated in KSHV (−) tumors are presented (Figure 4M). These include some examples of genes that were hypo-methylated and up-regulated during the transition from KSHV (+) cell to KSHV (+) tumor and are now hyper-methylated and down-regulated, such as Angpt4, Pdgfb, and Shank3. The negative regulators of the WNT signaling, APC and APC2, and Tle2 a transcriptional corepressor that binds to and Inhibits the transcriptional activation of CTNNB1 and TCF family members that mediate the Wnt signaling. This highlights the need to activate the Wnt signaling by other means, following the loss of the virus. Dow-regulation for the positive regulators of the epithelial/endothelial to mesenchymal transition, Zeb1 and Zeb2. In addition, regulators of RNA interference, Ago2, and regulators of chromatin organization, polycomb catalytic subunit Ezh1. Chromosome view of two genes with hyper-methylation in the regulatory region, clearly show increase in DNA methylation in the gene promoter (Figure 4N). Gene ontology analysis on STRING (https://string-db.org) for these hyper-methylated and down-regulated genes identified genes in the following cellular pathways (KEGG); PI3K-Akt signaling, Rap1 signaling, focal adhesion, ECM-receptor interaction, adrenergic signaling in cardiomyocytes, cholinergic and glutamatergic synapse (Figure 4O). Interestingly, pathways PI3K-Akt signaling, Rap1 signaling, axon guidance, microRNA in cancer, focal adhesion, and ECM-receptor interaction were upregulated and gene promoters were hypo-methylated during tumorigenesis with the virus, and these same pathways were down-regulated and gene promoters were hyper-methylated during tumor formation following loss of the virus (Figure 4G and 4O). This indicates that viral proteins/RNAs are necessary for both the hypo-methylation and up-regulation, but also for the maintenance of these pathways’ activation. These marks are eliminated following viral loss. We validated our global methylation analysis by performing qPCR with specific primers for several differentially methylated promoter regions (Figure S1).

### Analysis of the mutational landscape

KSHV-negative tumors belong to the same cell lineages as mECK36 tumors and share a significant overlap with the human KS transcriptome (Ma et al., 2013). Therefore, they are the best available control to be used in combination with KSHV-positive mECK36 tumors, which are a model of KSHV-dependent tumorigenesis (Mutlu et al., 2007), to assess KSHV-specific biology (Ma et al., 2013). This indicates that during *in vivo* tumorigenic growth host-cells acquire mutations that are able to convey tumorigenicity in the absence of KSHV. We have previously found one such mutation as KSHV (−) tumors and cells had a heterozygous D842V mutation in the tyrosine kinase (TK) domain of the PDGF receptor alpha (Cavallin et al., 2018). PDGFRA was wild-type in KSHV (+) tumors and cells, supporting the idea that PDGFRA is an oncogenic driver in KSHV tumors (Cavallin et al., 2018). We were able to find Skin KS biopsies in which only a small percentage of phospho-PDGFRA positive cells were LANA positive (Figure 5A). The existence of these KS lesions with low levels of KSHV infected cells suggests the existence of virus-independent hit-and-run mechanisms of sarcomagenesis whereby the KSHV oncovirus is able to induce irreversible genetic and epigenetic alterations. Interestingly, Figure 5B shows that KSHV (+) cells explanted from KSHV (+) tumors have much higher levels of ⍰H2AX phosphorylation than their cultured counterparts, which indicates that during the tumor growth KSHV (+) cells increased their foci of DNA repair. This suggests that during tumorigenic *in vivo* growth, there is an increase in DNA damage and; thus, KSHV (+) cells explanted from tumors are likely to have acquired irreversible host tumorigenic mutations that may contribute to overall tumor growth and may allow explanted cells to form tumors in the absence of KSHV. In order to analyze mutations acquired during *in vivo* tumorigenic growth we used the RNA-sequencing data and the Genome Analysis Toolkit (GATK) workflow to call for host mutations (Figure 5C). Detailed analysis of the mutational landscape of KSHV (+) and KSHV (−) cells and tumors was carried out taking in account the following criteria: 1) Transcripts <u>have to be expressed</u> in all compared samples (seq. depth >5). 2) We selected highly mutated genes: either one homozygous or two heterozygous mutations in various locations of the gene. One remarkable finding was that there were genes mutated in all six KSHV (+) tumors, but not in the KSHV (+) cell cultures (Figure 5D). Interestingly, 56 genes had recurrent mutations in the exact same position (87.5%). The only possible explanation for this finding is that these mutations were present already in the polyclonal population of KSHV (+) cells but were not detected by the RNA-sequencing analysis since they were present in a minimal percentage of cells. Yet, the consistent appearance of mutations in the same gene locations in all the tumors once the cells grew *in vivo* indicates that these mutations were enriched as they likely provided an *in vivo* survival advantage. Moreover, many of the mutations that “appear” in all KSHV (+) tumors “disappear” in explanted cells only to “reappear” when these KSHV (−) tumor cells are re-injected *in vivo* to form KSHV (−) tumors, further showing the reversibility of the selection process once cells are explanted and how selection operates again when these cells grow back *in vivo* (Figure 5D). Interestingly, network analysis of these mutations showed that many of them corresponded to innate immune genes including several of the TRIM family and the IFI family which are IFN inducible genes and a series of genes implicated in cytoplasmic DNA sensing (Figure 5E) and inflammasome/IFN activation. We found another very interesting set of mutations which appear de novo and manifested only on KSHV (−) tumor cells and KSHV (−) tumors. Clustering analysis of these mutations reveal a network comprising PDGFRA D842V, Pak1 (a Rac1 activated kinase) and Nucleolin mutations implicated in cell proliferation (Figure 5F). The analysis of these mutations points to KSHV tumorigenesis as a complex combination of effects from lytic KSHV oncogenesis combined with host mutations in innate immune genes that enable expression of those viral oncogenic genes and repress immunity against the virus allowing KSHV-dependent tumorigenesis to progress.

**FIGURE 5.**
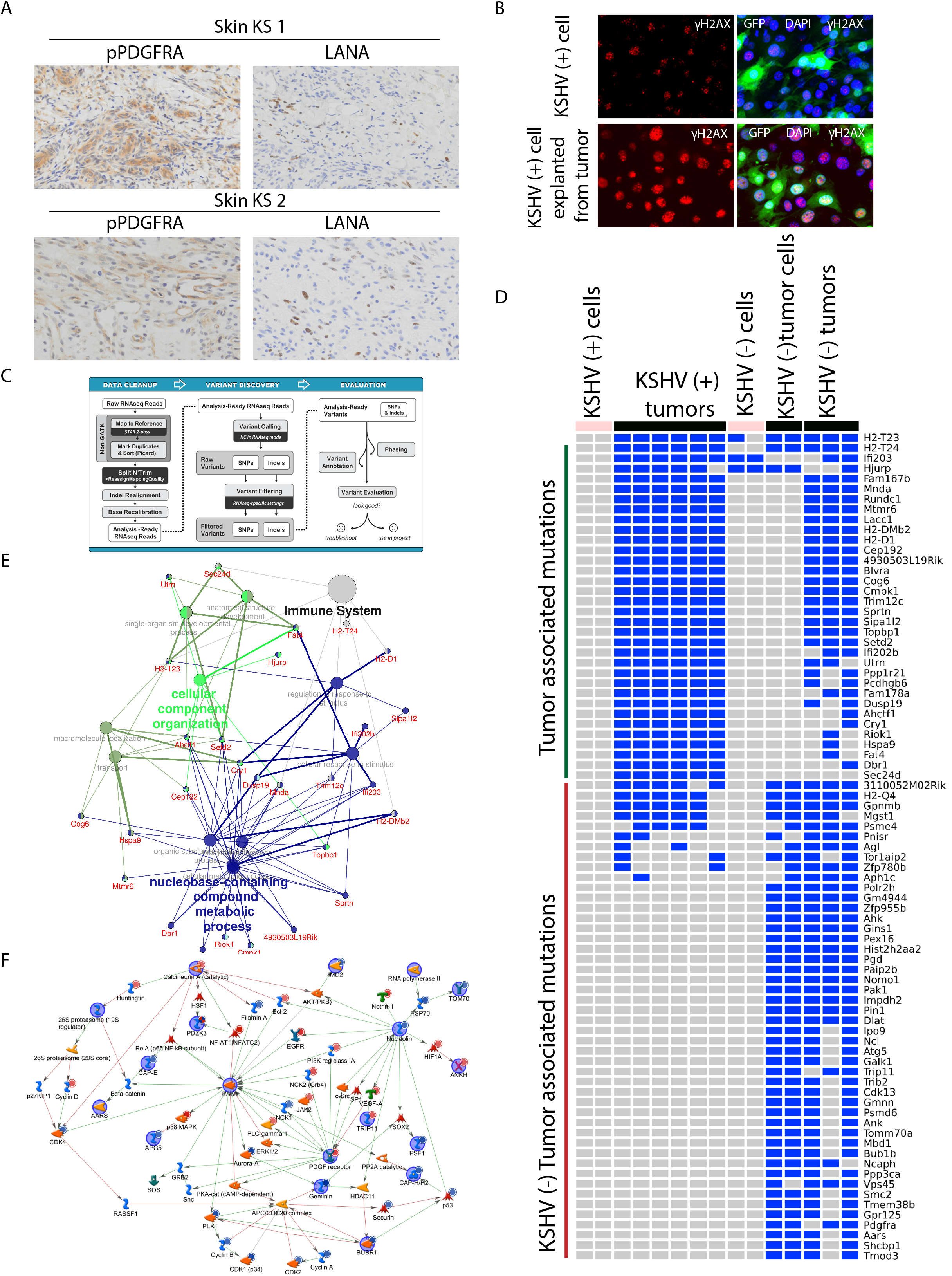
ANALYSIS OF THE MUTATIONAL LANDSCAPE. **(A)** Staining AIDS-KS biopsies from a ACSR tissue microarray (TMA) showing High phospho-PDGFRA and low LANA staining in two characteristic samples. **(B)** Immunofluorescence analysis of ⍰H_2_AX expression (red) was performed on GFP-positive KSHV (+) cells and GFP-positive KSHV (+) cells explanted from tumor; nuclei were counterstained with DAPI (blue). **(C)** The GATK workflow used to call host mutations. **(D)** Heatmap of the most frequently mutated genes. **(E)** Network analysis of mECK36 KSHV+ve tumors associated mutations **(F)** Network modeling with genes mutated in KSHV-ve mECK36 tumors, but not in mECK36 KSHV+ve cells or tumors.

## DISCUSSION

We have previously shown, by real-time qRT-PCR, that KSHV tumorigenesis in this mECK36 mouse KS-like model occurs with concomitant up-regulation of KSHV lytic genes and angiogenic ligands/receptors (Cavallin et al., 2018; Mutlu et al., 2007). Importantly, our present RNA-sequencing analysis also showed up-regulation of all the members of the PDGF family members in KSHV (+) tumors together with an increase in KSHV lytic gene expression. Moreover, pathway analysis of differential express genes (DEGs) showed that DNA methylation and Epigenetic regulation are the most relevant pathways involved in KSHV-dependent tumorigenesis (Figure 2A-F). This indicates that, as expected for a virus with strong epigenetic reprogramming capacities, KSHV-induced tumorigenesis is tightly linked to epigenetic regulation of host gene expression. This is also reinforced by the fact that KSHV-independent tumorigenesis occurred predominantly by Immune and Metabolic pathway regulation (Figure 2G-I). Interestingly, KSHV (+) tumors showed up-regulation of Immune cells infiltration and endothelial cell components when compared with KSHV (−) tumors by CIBERSORT analysis, which is consistent with the recruitment of inflammatory cells in the context of viral tumorigenesis and to the ability of KSHV to up-regulate angiogenesis respectively.

Our system allowed following whole genome CpG methylation of the host during tumorigenesis by KSHV. During tumorigenesis, we observed clear preference towards hypo-methylation in gene promoters. It is important to mention that the goal of this study was to identify the changes during tumorigenesis and not following infection, therefore the hyper-methylation immediately following infection is already present in the KSHV (+) cells. These results are in agreement with a previous study (Journo et al., 2018) that tried to address this question indirectly by comparing de-novo infection (in BJAB cells) to chronically infected PEL cells, and detected very similar hyper-methylation between de-novo and PEL, but dramatic difference towards hypo-methylation in PEL. While our study directly evaluates the methylation changes during transformation, both studies support the notion that hypo-methylation is the major epigenetic change during this process. Interestingly, hypo-methylation is also the major process during transformation of naïve B-cells following Epstein-Barr virus (EBV) infection (Hansen et al., 2014).

CpG DNA methylation is an epigenetic mark that can be faithfully maintained between generations (Mathieu et al., 2007; Wigler et al., 1981). Therefore, CpG methylation changes imposed by viral infection are expected to maintain following viral clearance. Our findings indicate that indeed some methylation changes are maintained following viral clearance, but some are lost, suggesting that the presence of viral encoded proteins or RNAs are necessary to maintain these epigenetic changes. In the case of KSHV (+) tumors these gene promoters are hypo-methylated and the genes are expressed, and in KSHV (−) tumors upon virus loss, these promoters becomes methylated again and repressed.

During the development of tumors following loss of KSHV we observed hyper-methylation of three main groups of genes: 1) genes that were previously hypo-methylated by the virus and viral gene products need to be expressed in order to keep these genes active, therefore viral loss leads to re-repression and hyper-methylation. 2) oncogenic pathways that are activated by the virus and are essential for tumor growth, upon viral loss the tumor cells need to find alternative ways to activate these pathways. Nice example is the Wnt signaling that is activated by KSHV encoded LANA (Fujimuro et al., 2003). During tumor growth without the virus we detected hyper-methylation of three repressors of the Wnt signaling, the tumor suppressors and the antagonists of the Wnt signaling, APC (Miyaki et al., 1993) and APC2 (van Es et al., 1999), and Tle2 a transcriptional corepressor that binds to and Inhibits the transcriptional activation of CTNNB1 and TCF family members that mediate the Wnt signaling (Brantjes et al., 2001). 3) changes due to cell differentiation/de-differentiation. KSHV was found to induce endothelial to mesenchymal (EdMT) transition (Cheng et al., 2011; Gasperini et al., 2012). In agreement with this transition, we observed hypo-methylation of ZEB1 during tumor development in KSHV infected cells. On the other hand, when tumors develop in the absence of the virus, we observed hyper-methylation of ZEB1 and ZEB2, that indicate a shift backwards mesenchymal to endothelial (MTEd) transition.

KSHV is a reprogramming virus encoding genes with powerful epigenetic regulatory abilities. Characterization of the CpG methylation footprint of KSHV infection showed a tendency towards hypo-methylation concomitant with KSHV tumorigenesis occurring along the lytic switch and hyper-methylation in comparing tumors infected versus un-infected with KSHV. The methylation level of the genome is controlled by two opposing activities; DNA methyltransferase, DNMT1, DNMT3a, and DNMT3b, and DNA demethylases, TET1, TET2 and TET3. Our analysis during tumor development of KSHV (+) tumors detected promoter hypo-methylation and up-regulation of the demethylase Tet2 gene. This observation can provide a mechanistic explanation for the hypo-methylation that occurs during tumorigenesis. Moreover, we found PDGF signaling members (PDGFA, PDGFB and PDGFRA) also hypo-methylated and up-regulated in these transitions as well, further illustrating another KSHV-dependent mechanism to maintain the activation of this oncogenic signaling pathway. These results showed the importance of DNA methylation regulation in the process of KSHV-dependent oncogenesis.

The existence of KS lesions with varying levels of KSHV infected cells suggests also the existence of virus-independent hit-and-run mechanisms of sarcomagenesis whereby the KSHV oncovirus is able to induce irreversible genetic and epigenetic alterations. This is also supported by the fact that KSHV is a reprogramming virus and the occurrence of host oncogenic alterations in KS lesions (Nicolaides et al., 1994; Pillay et al., 1999). We previously found that KS lesions overexpress Rac1 and that oxidative stress plays a major role in KS sarcomagenesis (Ma et al., 2013; Ma et al., 2009). ROS mediated *in vivo* genetic damage could lead to accumulation of mutations that could contribute to the viral oncogenesis process and may compensate for the loss of virus as found in some KS lesions (Figure 5A). We found that cells explanted from tumors display much more foci of DNA repair which is consistent with increased “in tumor” DNA damage and DDR (Figure 5B).

In analyzing the mutational landscape we made an astonishing observation, there were genes mutated in all six KSHV (+) tumors, but not in the KSHV (+) cell cultures (Figure 5D). This indicates that in the context of KSHV tumorigenesis not only selects for the presence of the virus (Mutlu et al., 2007) but also pre-existing host mutations that --as the KSHV episome--only provide a selective advantage *in vivo*. Interestingly many of these mutations appear to be in genes regulating viral DNA and innate immunity genes. Some of these TRIM and IFI family genes, which are IFN inducible genes, have already shown to be important for KSHV lytic reactivation (Fukushi et al., 2003; Full et al., 2019; Roy et al., 2016). It is likely that these mutations allow the KSHV oncovirus to express the oncogenic lytic program and that they allow for a permissive environment of inflammatory and viral tumorigenesis, which occurs in the context of DDR that may be impeded by innate immune DNA sensors.

We found that KSHV oncogenesis also induced the accumulation of de novo mutations that were not present in KSHV (+) cells and tumors. These mutations were first selected upon KSHV loss *in vitro* and would be responsible—together with PDGFRA D842V-- to be driving tumorigenesis compensating KSHV loss (Figure 5D). We recently found that the most prominent of these mutations is the one activating the KS oncogenic driver PDGFRA (PDGFRAD842V), that allows the cell to maintain PDGF driven tumorigenesis in the absence of KSHV (Cavallin et al., 2018). Interestingly, this D842V mutation is the most common activating mutation of PDGFRA found in GIST. Network analysis of the other de novo mutations point to the existence of a PDGFRA-Rac1 driven network that is consistent with the Rac1 overexpression in AIDS-KS (Ma et al., 2009) and mECK36 KSHV (+) and KSHV (−) tumors as well as the proposed role of ROS in AIDS-KS as shown by mECK36 NAC sensitivity (Ma et al., 2013).

## Materials and methods

### Cell Culture and Reagents

mECK36 cells KSHV (+) were obtained and cultured as previously described (Mutlu et al., 2007). KSHV (−) tumor cells were obtained from mECK36 tumor explants after Bac36 episome loss achieved by growth without hygromycin selection as previously described (Mutlu et al., 2007).

### RNA-Seq analysis

RNA was isolated and purified using the RNeasy mini kit (Qiagen). RNA concentration and integrity were measured on an Agilent 2100 Bioanalyzer (Agilent Technologies). Only RNA samples with RNA integrity values (RIN) over 8.0 were considered for subsequent analysis. mRNA from cell lines and tumor samples were processed for directional mRNA-seq library construction using the Preparation Kit according to the manufacturer’s protocol. We performed paired-end sequencing using an Illumina NextSeq500 platform. The short sequenced reads were mapped to the mouse reference genome (GRCm38.82) by the splice junction aligner TopHat V2.1.0. We employed several R/Bioconductor packages to accurately calculate the gene expression abundance at the whole-genome level using the aligned records (BAM files) and to identify differentially expressed genes between cell lines and cell lines and tumors. Briefly, the number of reads mapped to each gene based on the TxDb. Mmusculus gene ensembls were counted, reported and annotated using the Rsamtools, GenomicFeatures, GenomicAlignments packages. To identify differentially expressed genes between cell lines and tumor samples, we utilized the DESeq2 test based on the normalized number of counts mapped to each gene. Functional enrichment analyses were performed using the ClueGo Cytoscape’s plug-in (http://www.cytoscape.org/) and the InnateDB resource (http://www.innatedb.com/) based on the list of deregulated transcripts. Data integration and visualization of differentially expressed transcripts were done with R/Bioconductor.

### Methylated DNA Binding Protein sequencing (MBD-seq)

#### Methylated DNA Enrichment

Genomic DNA was isolated using DNeasy Blood & Tissue Kit (QIAGEN) and sheared by sonication to fragments of ~500bp. The sheared DNA (1μg) was added to 10 μl MBD-Bead slurry (MethylMiner DNA Enrichment Kit, Invitrogen, Carlsbad, CA) and incubated on a rotating mixer for 1 hr, as described previously (Shamay et al., 2012b). The DNA fragments were eluted into distinct subpopulations based on the degree of methylation; non-captured fraction (NC, representing un-methylated DNA fragments), 450 mM NaCl (representing partially methylated DNA fragments) and 2000 mM NaCl (representing methylated DNA fragments). The fractions were then ethanol precipitated and re-suspended in H_2_O.

#### DNA sequencing and analysis

MBD2 enriched DNA fractions were subjected to library preparation with NEBNext^®^ Ultra^™^ || DNA library prep kit for Illumina (NEB #E7645) at the next generation genomic center in the Azrieli Faculty of Medicine, Bar-Ilan University. The DNA libraries were sequenced on Illumina HiSeq2000, with HiSeq rapid 100PE. Data analysis was performed on the Partek flow platform, raw reads were aligned to the mouse genome mm10 (GRCm38/mm10) with BWA and more than 80% reads were uniquely aligned to the reference genome (Kim et al., 2013)(Kim et al., 2013)(Kim et al., 2013)(Kim et al., 2013)(Kim et al., 2013)(Kim et al., 2013)(Kim et al., 2013). Mapped reads were analyzed with MACS2 to generate peaks and annotate differentially methylated regions. Peaks with False discovery rate (FDR) ≤5% & Fold change (FC) ≥2, were considered differentially methylated. When peaks were located between1500 bp upstream and 200 bp downstream of the transcription start site (TSS), they were included in the differentially methylated gene promoters.

### Immunofluorescence Staining

Immunofluorescence assay (IFA) was performed as previously described (Mutlu et al., 2007). Cells were fixed in 4% paraformaldehyde for 10 min and washed with PBS. Cells were o permeabilized in 0.2% Triton-X/PBS for 20 min at 4°C. After blocking with 3% of BSA in PBS and 0.1% Tween 20 for 60 min, samples were incubated with Primary antibodies overnight at 4C. After PBS washing, samples were incubated with fluorescent secondary antibodies for 1 hour (Molecular Probes), washed and mounted with ProLong Gold antifade reagent with DAPI (Molecular Probes). Images were taken using a Zeiss ApoTome Axiovert 200M microscope.

### Animal Studies

All mice were housed under pathogen-free conditions. Tumor studies were done in 4- to 6-week-old nude mice obtained from the National Cancer Institute. Tumors were generated 5 by subcutaneous injection of mECK36 cells (3 × 10^5^ cells) as previously described (Mutlu et al., 2007).

### Clinical tissue analysis

Skin KS biopsies were analyzed from an ACSR (The AIDS and Cancer Specimen Resource) tissue microarray. Immunohistochemistry of clinical tissue microarrays was performed using a standard protocol of the Immunohistochemistry Laboratory of the Department of Pathology at the University of Miami. Antibody staining of p-PDGFRA from R&D Systems (Minneapolis, MN) was diluted to 1:30 and LANA from Abcam (Cambridge, MA) was diluted 1:40.

### Ethics Statement

The animal experiments have been performed under UM IACUC approval number 13-093. The University of Miami has an Animal Welfare Assurance on file with the Office of Laboratory Animal Welfare (OLAW), National Institutes of Health. Additionally, UM is registered with USDA APHIS. The Council on Accreditation of the Association for Assessment and Accreditation of Laboratory Animal Care (AAALAC International) has continued the University of Miami’s full accreditation.

## Acknowledgments

We would like to thank the Azrieli Faculty of Medicine Genomics Center for MBD2-sequencing, and the BCF Technion Genomics Center for methylation data analysis. We are grateful for the support of the Elias, Genevieve and Georgianna Atol Charitable Trust to the Daniella Lee Casper Laboratory in Viral Oncology. This work was supported by grants from the Israel Science Foundation (https://www.isf.org.il) to M.S. (1134/16), and Research Career Development Award from the Israel Cancer Research Fund (https://www.icrfonline.org/) to M.S. (01282). The funders had no role in study design, data collection and analysis, decision to publish, or preparation of the manuscript.

