## Supplementary Tables 1 and 2 for "RNA-sequencing analysis of a multistep and hit-and-run cell and animal model of KSHV tumorigenesis reveal the roles of mutations, CpG methylation, and viral-infection footprints in oncogenesis"

| | Hyper | Hypo |
|---|---|---|
| **Biological process** | no terms | pancreatic A cell differentiation<br>signal transduction involved in regulation of gene expression<br>establishment of endothelial barrier regulation of cell shape<br>astrocyte differentiation<br>lacrimal gland development response to cAMP<br>positive regulation of dendritic spine development<br>response to organophosphorus<br>response to purine-containing compound<br>negative regulation of female gonad development<br>glial cell fate commitment<br>platelet-derived growth factor receptor-beta signaling pathway<br>regulation of reactive oxygen species metabolic process<br>enteroendocrine cell differentiation<br>positive regulation of reactive oxygen species metabolic process<br>positive regulation of smooth muscle contraction<br>embryonic eye morphogenesis<br>embryonic camera-type eye morphogenesis<br>response to amine stimulus<br>nuclear chromatin<br>ESC/E(Z) complex<br>pical dendrite |
| **Pathway** | no terms | Genes involved in Regulation of Insulin Secretion<br>Maturity onset diabetes of the young<br>Genes involved in Regulation of gene expression in beta cells<br>Genes involved in Integrin cell surface interactions<br>Genes involved in PECAM1 interactions<br>Genes involved in Regulation of beta-cell development<br>Genes involved in Signal regulatory protein (SIRP) family interactions<br>Genes involved in Regulation of KIT signaling Regulation of p38-alpha and p38-beta |
| **Oncogenic signature** | no terms | Genes up-regulated in Sez-4 cells (T lymphocyte) that were stimulated with IL2.<br>Genes up-regulated in TIG3 cells (fibroblasts) upon knockdown of EED gene.<br>Immune or inflammatory genes induced by NF- kappaB in primary keratinocytes and fibroblasts.<br>Genes defining the KRAS dependency signature.<br>Genes up-regulated in retina cells from CRX and NRL double knockout mice,<br>Genes down-regulated in retina cells from NRL knockout mice.. |
| **Predicted promoter motifs** | no terms | Motif CACGTGS matches MYC: v-myc myelocytomatosis viral oncogene homolog (avian)<br>Motif GGGGAGGG matches MAZ: MYC- associated zinc finger protein<br>Motif NNGTNRCNATRGYAACNN matches RFX1: regulatory factor X, 1 (HLA class II expression)<br>Motif TGASTMAGC matches NFE2: nuclear factor (erythroid-derived 2), 45kDa<br>Motif CARAACTAGGNCAAAGGTCA matches PPARA: peroxisome proliferative activated receptor, alpha.<br>Motif GCACCCAWGGGTGM matches ZNF423: zinc finger protein 423 |

**Table S1: Biological processes and pathways identified in GREAT during the transition from KSHV (+) cells to KSHV (+) tumors.**

| | Hyper | Hypo |
|---|---|---|
| **Biological process** | actin filament organization<br>regulation of cell shape<br>cell-substrate adhesion<br>actin filament bundle assembly<br>phagocytosis<br>regulation of phagocytosis<br>micturition<br>alanine catabolic process<br>cell-matrix adhesion<br>smooth muscle contraction involved in micturition<br>Rac protein signal transduction<br>epithelial cell differentiation involved in kidney development<br>urinary tract smooth muscle contraction<br>mannosylation | no terms |
| **Pathway** | Members of the BCR signaling pathway<br>Small cell lung cancer<br>RXR and RAR heterodimerization with other nuclear receptor<br>Retinoic acid receptors-mediated signaling | no terms |
| **Oncogenic signature** | cancer neighborhood for:<br>SELL<br>MYD88<br>RAGE<br>LTK<br>PML | no terms |
| **Predicted promoter motifs** | no terms | Motif CACSCCA matches SREBF1 |

**Table S2 : Biological processes and pathways identified in GREAT during the transition from KSHV (+) tumors to KSHV (-) tumors.**
