## Supplementary figures and images for "RNA-sequencing analysis of a multistep and hit-and-run cell and animal model of KSHV tumorigenesis reveal the roles of mutations, CpG methylation, and viral-infection footprints in oncogenesis"

### Supplementary Figure 1

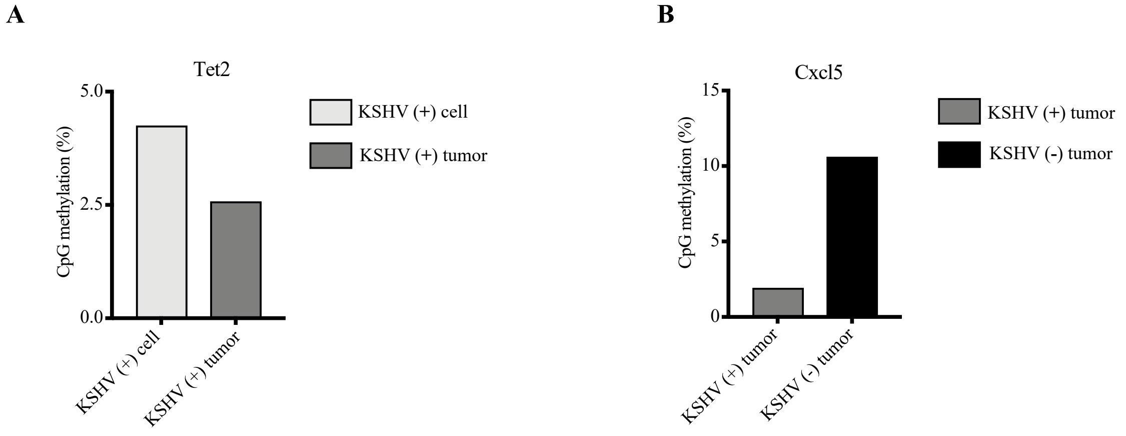
